# Comparison of the Drug Entrapment Efficiency of Almond Gum (Badam Pisin) to Span-60 Niosomes by Folin-Ciocalteu Assay

**DOI:** 10.1101/2021.04.03.438312

**Authors:** R Hari Krishnan

## Abstract

Almond gum (Badam Gum/ Badam Pisin) is a natural polymer composed of mainly carbohydrates (>90%) which makes it a potential candidate in targeted drug delivery. It contains 46% L-arabinose which is known to have fluid entrapment capacity. Various methods of drug targeting against tumor have been developed but due to poor solubility in body fluids, the bioavailability of the drug is reduced and so is its effectiveness. Almond gum is known to be water-soluble so the required bioavailability will be achieved and the drug dosage will also be minimal. Since it is of natural origin side effects will be less compared to that of artificially prepared drugs. Studies show that almond gum is known to have various health benefits including increasing immunity and acting as a coolant. *Terminalia catappa* (Indian almond), the source of almond gum is abundant in the Indian subcontinent hence making it more economical than chemical surfactants. The objective of this study is to determine whether Almond gum has a higher drug entrapment efficiency than that of Niosomes.

## Introduction

Conventional drug administration is associated with various drawbacks including limited effectiveness, poor bioavailability, lack of sensitivity, and increased toxicity. To combat these drawbacks nanocarrier mediated drug carriers have been evolving recently which effectively targets specific diseased tissues of the body, avoiding any interaction with healthy tissues. This type of drug delivery system is called Targeted Drug Delivery whose goal is to have a localized and protected drug interaction with the affected tissue over a prolonged period.

Almond gum extracted from *Terminalia catappa* is a naturally occurring polymer composed of *carbohydrates* (92.36%), proteins (2.45%), and trace amounts of fats (0.85%). The drug entrapping capacity of Almond gum arises from its high concentrations of L-arabinose (46.83%) which is known to have a high fluid retention capacity. The other carbohydrate components present in Almond gum are galactose (35.49%) and uronic acid (5.97%) with traces of rhamnose, mannose, and glucose. Coming from a natural origin with added health benefits and being highly economical Almond gum as a drug carrier can significantly reduce various drawbacks of chemically synthesized Niosomes.

In this report, we compare the drug entrapment efficiency of commercially available Almond gum with chemically synthesized Span-60-cholesterol Niosomes. The drug used in this study are polyphenols extracted from commercially available green tea leaves. The total concentration of the drug administered for entrapment was 10mg/ml. The total polyphenol concentration that is not entrapped in the vesicles and present in the supernatant were measured using Folin–Ciocalteu (F–C) assay for identification of phenols wherein the unknown concentration of non-entrapped drug can be identified by comparing its absorbance (at 765nm) with its corresponding concentration using a gallic acid standard plot.

The drug entrapment efficiency is calculated for both Almond gum and niosomes using the formula-

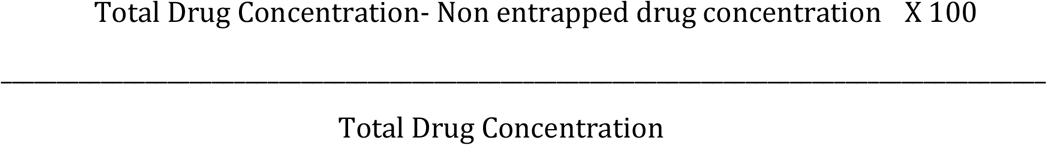

The sample with a higher drug entrapment efficiency is associated with higher bioavailability. It was found that Almond gum had a higher entrapment efficiency than the Niosomes hence explaining its importance in modern drug delivery systems.

## Materials required

1. For polyphenol extraction-Green tea leaves dried, ethanol, funnel, conical flask, filter paper.
2. For Niosome preparation-Span 60, cholesterol, chloroform, weighing balance, round bottom flasks, rotary evaporator, centrifuge, glasswares (beakers, measuring cylinders)
3. Test for phenolic group-Fe (III) chloride, polyphenol extract
4. Almond gum vesicle preparation-Powdered almond gum, ethanol, distilled water, magnetic stirrer with heater, centrifuge, falcon tubes, polyphenol extract
5. Polyphenol estimation by Folin–Ciocalteu reagent-Sodium bicarbonate, Folin-Ciocalteu reagent, distilled water, test tubes, pipettes, phenol extract (for standard), drug entrapped vesicles (both noisome and almond gum)

## Methods

1. **Polyphenol extraction-** The drug used for entrapment are polyphenols and green tea is a good source of this antioxidant. 100 ml of distilled water is heated in separate flasks, and 1g of dried green tea leaves are added to each of the flasks and are heated for 30min. This mixture is then filtered to remove the leaves and the filtered solution now contains polyphenols. Confirmation of polyphenol is done by Fe (III) test wherein phenol reacts with *FeCl_3_* to produce a Fe containing violet complex.

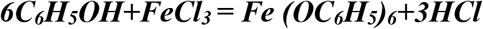 ***Fe (OC_6_H_5_)_6_*** gives a characteristic dark blue/purple colour indicating the presence of phenols.
2. **Preparation of Niosomes-** For the preparation of niosomes, Span-60 surfactant and cholesterol (powder) stocks were made and dissolved in chloroform in a 1:1 ratio. Chloroform was used because it is a fat-soluble solvent. This mixture was then placed in a rotary vacuum evaporator at 40^0^C until the liquid was evaporated and a thin translucent lipid layer was visible at the bottom of the flask. To this 5ml of the extracted polyphenol solution was added and incubated for a day. Hydration of the thin layer causes the hydrophobic end of the niosome to curl around and form a spherical vesicle entrapping the drug. This is then centrifuged for 5min at 10000rpm to precipitate the pellet and supernatant. 100ul of this supernatant is carefully extracted and subjected to F-C assay to find the concentration of the non-entrapped drug.
3. **Almond gum entrapment-** Commercially available almond gum, also known as Badam Pisin was blended into powder using a mixer. The powder was then dissolved in 10ml of distilled water and heated in a magnetic stirrer for the powder to be dissolved. Once the powder is dissolved, 5ml of the extracted polyphenol (same volume that was added for niosome) is then added to this solution and is incubated for a day at room temperature. After incubation, this mixture was centrifuged for 5min at 10000rpm to obtain the pellet and supernatant. 100ul of this supernatant was extracted and is subjected to F-C assay to estimate the concentration of the non-entrapped drug.
4. **Polyphenol estimation by Folin–Ciocalteu assay-** The Folin–Ciocalteu reagent is a mixture of tungstates and molybdates that works on the mechanism of an oxidation-reduction reaction. The method strongly relies on the reduction of the mixture heteropolyphosphotungsates–molybdates by the phenolic compound which results in the formation of blue-colored chromogen. Hence more the absorption value more is the drug present in the supernatant and less is the intake by the vesicle. The procedure involves the preparation of a 7.5% solution of sodium bicarbonate responsible for providing the required alkaline medium. 100ul of the almond gum supernatant is taken and 900ul of water is added to make the total volume as 1ml. To which 1ml sodium bicarbonate and 1ml F-C reagent are added to make the total volume of the mixture be 3ml. The same procedure is repeated with 100ul niosome supernatant. These mixtures are then subjected to a spectrophotometer to measure their absorbance. The unknown concentration of the drug in the supernatant can be estimated by plotting the gallic acid standard curve by preparing a gallic acid stock solution of different concentrations and measuring its absorbance. This is called the Gallic acid equivalence method (GAE).

## Results and Discussion-

The study of drug retention kinetics was performed between natural polymer Almond gum and chemically synthesized Span-60 Niosomes. Readings from the F-C assay showed that the entrapment efficiency of natural almond gum is higher when compared to the niosome. Plotting a standard graph between concentration and Optical density using gallic acid standards gives the concentration of drug present in the supernatant (non-entrapped). Subtracting this value with the total drug concentration we can get the concentration of polyphenol encapsulated in almond gum and niosomes respectively.

100uL volume of the supernatants of almond gum and niosomes were subjected to phenol estimation test using F-C reagent and the following absorption values were obtained-

Fig.1 shows the standard plot of gallic acid curve from the data of Table 2 where, concentration is taken in the X-axis and optical density is taken in the Y-axis. The unknown concentration of the drugs in the almond gum and niosome supernatants was found by plotting the X-intercept corresponding to the OD values respectively.

**Fig.1 –.**
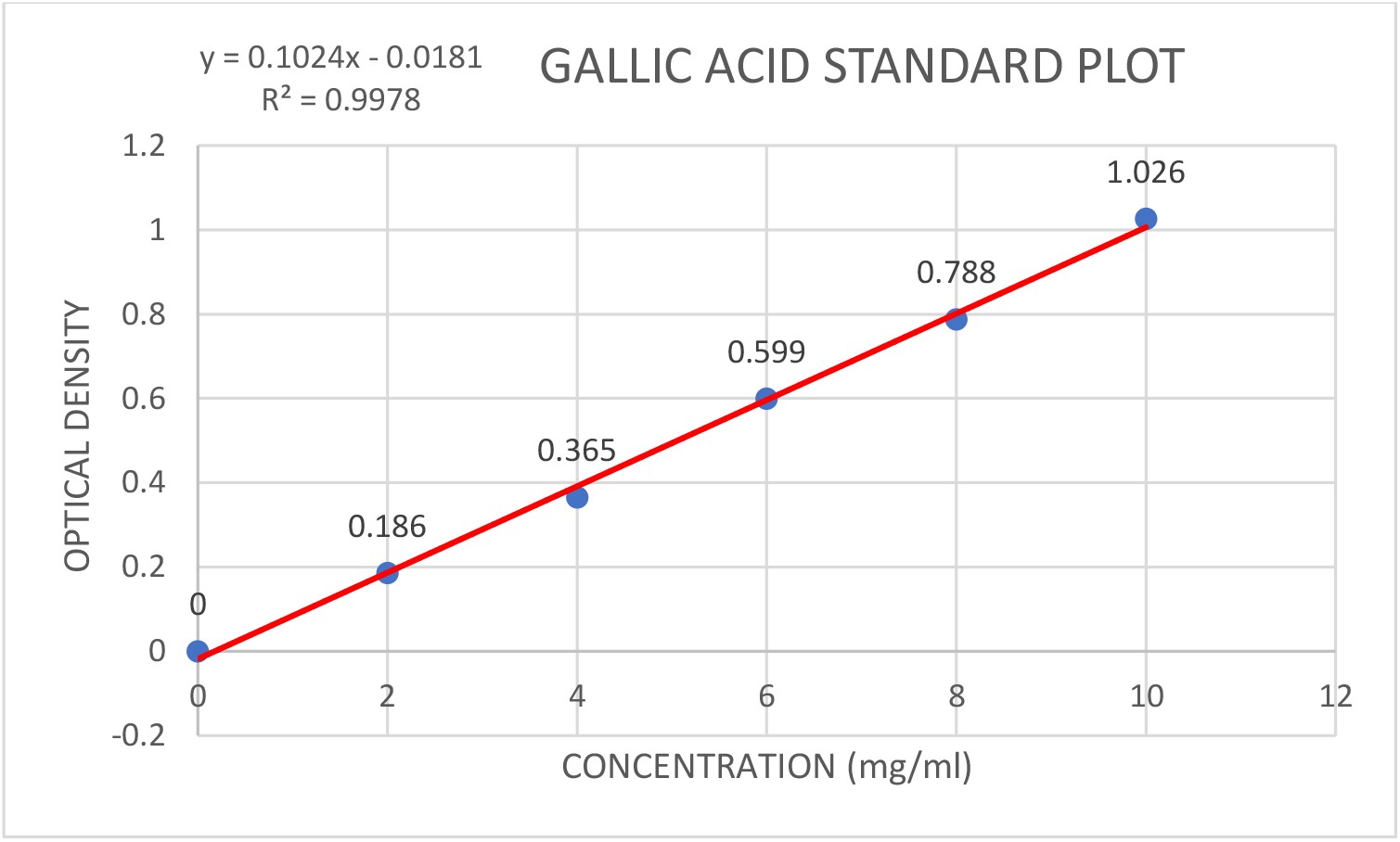
Gallic acid standard plot for identification of the unknown concentration of drug. Concentration is taken on the X-axis and Optical density is taken in the Y-Axis, the x intercept for the known OD is calculated using the line equation mentioned within the figure.

**Table 1-.**
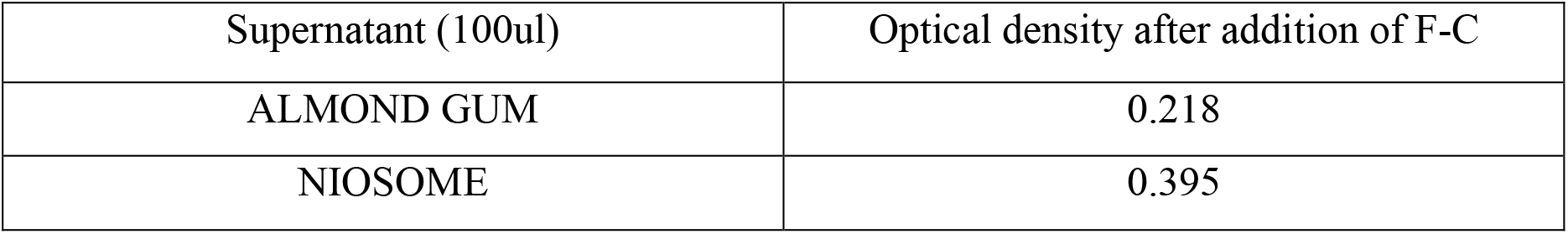
Optical density of supernatants after addition of F-C

**Table 2-.** Gallic acid standard values

| Concentration of Gallic acid (mg/ml) | Optical density |
| --- | --- |
| 2 | 0.186 |
| 4 | 0.365 |
| 6 | 0.599 |
| 8 | 0.788 |
| 10 | 1.026 |

Solving the line equation of the standard plot in Fig.1, we get x (concentration) of drug in almond gum supernatant = 2.305mg/ml and x (concentration) of drug in niosome supernatant = 4.034 mg/ml. Total Drug concentration = 10mg/ml

The amount of drug entrapped in almond gum = 10-2.305 = 7.695mg/ml; The amount of drug entrapped in niosome vesicle = 10-4.034 = 5.966 mg/ml.

The drug entrapment efficiency of Almond Gum=

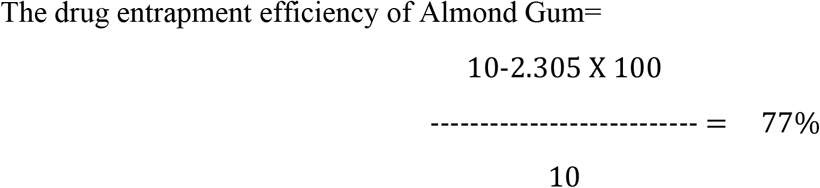

The drug entrapment efficiency of Niosome =

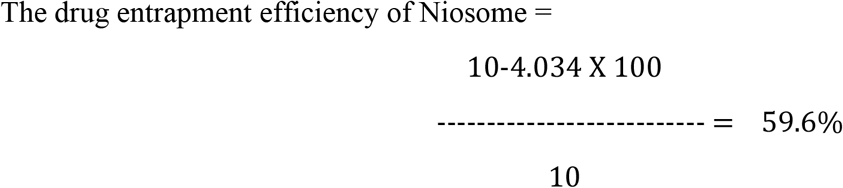

## Conclusion

In conclusion, our *in-vitro* drug entrapment study has shown that the natural polymer almond gum has a significant drug entrapment capacity. There is still a lot of work to be done on almond gum to precisely find out whether it is an ideal drug carrier. Various pharmacodynamic, pharmacokinetics, and *in-vivo* studies in the future might be beneficial in understanding more about the drug entrapping and releasing nature of Almond gum. There has been a continued demand for novel natural biomaterials or biopolymers for their quality of being biodegradable, biocompatible, readily available, renewable, and low toxicity. Beyond identifying such polysaccharides and proteins natural biopolymers, research on making them more stable under industrial processing environment and biological matrix through techniques such as crosslinking are among the most advanced research area nowadays.

